# SLIC-CAGE: high-resolution transcription start site mapping using nanogram-levels of total RNA

**DOI:** 10.1101/368795

**Authors:** Nevena Cvetesic, Harry G. Leitch, Malgorzata Borkowska, Ferenc Müller, Piero Carninci, Petra Hajkova, Boris Lenhard

## Abstract

Cap analysis of gene expression (CAGE) is a methodology for genome-wide quantitative mapping of mRNA 5’ends to precisely capture transcription start sites at a single nucleotide resolution. In combination with high-throughput sequencing, CAGE has revolutionized our understanding of rules of transcription initiation, led to discovery of new core promoter sequence features and discovered transcription initiation at enhancers genome-wide. The biggest limitation of CAGE is that even the most recently improved version (nAnT-iCAGE) still requires large amounts of total cellular RNA (5 micrograms), preventing its application to scarce biological samples such as those from early embryonic development or rare cell types. Here, we present SLIC-CAGE, a Super-Low Input Carrier-CAGE approach to capture 5’ends of RNA polymerase II transcripts from as little as 5-10 ng of total RNA. The dramatic increase in sensitivity is achieved by specially designed, selectively degradable carrier RNA. We demonstrate the ability of SLIC-CAGE to generate data for genome-wide promoterome with 1000-fold less material than required by existing CAGE methods by generating a complex, high quality library from mouse embryonic day (E) 11.5 primordial germ cells.

## INTRODUCTION

Cap analysis of gene expression (CAGE) is used for genome-wide quantitative identification of polymerase II transcription start sites (TSSs) at a single nucleotide resolution (Shiraki et al. 2003) as well as 5’end-centred expression profiling of RNA polymerase II (RNAPII) transcripts. The region surrounding a TSS (approximately 40 bp upstream and downstream) represents the core promoter, where the transcription initiation machinery and general transcription factors bind to direct initiation by RNAPII (Smale and Kadonaga 2003). Information on exact TSS positions in the genome improves identification of core promoter sequences and led to the discovery of new core promoter and active enhancer sequences (Andersson et al. 2014; Consortium et al. 2014; Haberle et al. 2014) (reviewed in Lenhard et al (Lenhard et al. 2012) and Haberle et al (Haberle and Lenhard 2016)). The current knowledge of core promoter sequences identified by CAGE has uncovered their regulatory role on an unprecedented scale. CAGE detected TSS profiles represent an accurate and quantitative readout of promoter utilisation, their patterns reflect ontogenic, cell type specific and cellular homeostasis-associated dynamic profiles which allows promoter classification and informs about the diversity of promoter level regulation. This has led to increased use of CAGE techniques and their application in high profile research projects like ENCODE (Consortium 2012), modENCODE (Celniker et al. 2009), FANTOM3, and FANTOM5 (Kawaji et al. 2017). Finally, CAGE is proving to be invaluable for uncovering disease-associated novel TSSs (Boyd et al. 2018) that can be used as diagnostic markers, for associating effects of GWAS-identified loci with TSSs (Blauwendraat et al. 2016; Cusanovich et al. 2016), and for facilitating design of CRISPRi experiments (Qi et al. 2013).

Central to CAGE methodology is the positive selection of RNA polymerase II transcripts using the cap-trapper technology (Carninci et al. 1996). This technology uses sodium periodate to selectively oxidize vicinal ribose diols present in the cap structure of mature mRNA transcripts, facilitating their subsequent biotinylation. RNA is first reversely transcribed using a random primer (N_6_TCT) and converted to RNA:cDNA hybrids, followed by oxidation, biotinylation and treatment with RNase I to select only full-length RNA:cDNA hybrids; *i.e.* cDNA that has reached the 5’end of capped mRNA during reverse transcription will protect RNA against digestion with RNase I. Purification of biotinylated RNA:cDNA hybrids is then performed using streptavidin-coated paramagnetic beads. These steps ensure that incompletely synthesized cDNA and cDNA synthesized from uncapped RNAs are eliminated from the sample. The initial CAGE protocol required large amounts of starting material (30-50 μg of total cellular RNA) and used restriction enzyme digestion to generate short reads (20 bp) (Kodzius et al. 2006), whereas the later versions reduced the starting amount 10-fold and generated slightly longer reads (27 bp) with increased mappability (Takahashi et al. 2012).

The latest CAGE protocol using cap-trapping is nAnT-iCAGE (Murata et al. 2014) and it is the most unbiased method for genome-wide identification of TSSs. It excludes PCR amplification as well as restriction enzymes used to produce short reads in previous CAGE versions. However, at least 5 μg of total RNA material is still required for nAnT-iCAGE. To address this, an alternative, biochemically unrelated approach, nanoCAGE, was developed for samples of limited material availability (50-500 ng of total cellular RNA) (Plessy et al. 2010; Poulain et al. 2017). NanoCAGE uses template switching (Zhu et al. 2002) instead of the cap-trapper technology to lower the starting material. Template switching is based on reverse transcriptase’s ability to add extra cytosines complementary to the cap, which are then used for hybridization of the riboguanosine-tailed template switching oligonucleotide to extend and barcode only the 5’ full length cDNAs. Despite its simplicity, nanoCAGE has limitations that make it inferior to classic CAGE protocols: 1) template switching has been shown to be sequence dependent and therefore biased (Tang et al. 2013), potentially compromising the determination of preferred TSS positions; 2) production of libraries from 50 ng of total RNA often requires 20-35 PCR amplification cycles, leading to low-complexity libraries with high levels of duplicates. Although nanoCAGE methodology implements unique molecular identifiers (UMIs) (Kivioja et al. 2011; Poulain et al. 2017), their use for removal of PCR duplicates is often complicated due to problems with achieving truly randomly synthesised UMI’s and errors in sequencing (Smith et al. 2017).

Despite improvements in the CAGE methodology, the amount of input RNA needed for unbiased genome-wide identification of TSSs constitutes a true limitation when cells, and therefore RNA, are difficult to obtain. This is the case when working with embryonic tissue or early embryonic stages, rare cells types, FACS sorted selected cells, heterogeneous tumours, or diagnostic biopsies. Here, we present SLIC-CAGE, a Super-Low Input Carrier-CAGE approach that is based on cap-trapper technology and can generate unbiased high-complexity libraries from 5-10 ng of total RNA. Thus far the cap-trapper step has been the limiting factor in the reduction of the amount of required starting material. To facilitate the cap-trapper technology on the nanogram scale, representing capped RNA from as low as hundreds of eukaryotic cells, samples of the total RNA of interest are supplemented with novel pre-designed carrier RNAs. Prior to sequencing, the carrier is efficiently removed from the final library using homing endonucleases that target recognition sites embedded within the sequences of the carrier molecules, leaving only the target mRNA library to be amplified and sequenced. The specificity and the long recognition motifs of homing endonucleases ensure that no DNA derived from sample RNA is degraded in the process.

We have tested and validated SLIC-CAGE on a wide-range of starting material amounts (1-100 ng of total RNA) from *Saccharomyces cerevisiae* and *Mus musculus* using the current nAnT-iCAGE protocol, as a gold standard. Additional direct comparison between SLIC-CAGE and the latest nanoCAGE protocol (Poulain et al. 2017) showed that SLIC-CAGE strongly outperforms nanoCAGE in sensitivity, resolution and reproducibility. SLIC-CAGE produced unbiased libraries of higher complexity and quality than nanoCAGE, even when constructed using low total RNA input (5-10 ng compared to 500 ng). Finally, we demonstrate that SLIC-CAGE enables reliable genome-wide promoter-centric biological discovery and promoter classification using as little as 5-10 ng of total RNA material and validate its applicability by generating a high quality TSS landscape of mouse primordial germ cells (PGCs) embryonic (E) day 11.5.

## RESULTS

### Development of SLIC-CAGE

In typical CAGE protocols, the cap-trapper step needs at least 5 μg of total RNA (Murata et al. 2014) and is therefore the limiting factor. This step has been difficult to scale down as it involves the pull-down of biotinylated capped RNA using streptavidin beads. In such situations, a common biochemical approach to prevent sample loss is the use of carriers; *i.e.* inert non-interfering molecules to minimize sample loss caused by nonspecific adsorption and to improve specificity in affinity purification steps. However, unless there is a way to selectively remove the carrier afterwards, the carrier signal will dominate the sequenced sample and therefore lead to orders of magnitude of reduced sequencing depth of the sample itself.

To solve this problem and enable profiling of minute amounts of RNA, we set out to design a carrier RNA that will be similar in size distribution and the percentage of capped RNA to the cellular RNA, but whose cDNA will be possible to selectively degrade without affecting the cDNA originating from the sample.

We constructed the synthetic gene used as a template for run-off *in vitro* transcription of the carrier RNA (Supplemental Fig. S1A,B, Supplemental Table S1, see Methods for details). The synthetic gene is based on the *Escherichia coli* leucyl-tRNA synthetase sequence for two main reasons. First, we wanted to avoid mapping to eukaryotic genomes. Secondly, leucyl-tRNA synthetase is a housekeeping gene from a mesophilic species and therefore its sequence is not expected to form strong secondary structures that would reduce its translation *in vivo*, or reduce the efficiency of reverse transcription to form RNA:cDNA hybrids. We made this carrier selectively degradable by embedding it with multiple recognition sites of two homing endonucleases, I-CeuI and I-SceI (Fig. 1A, Supplemental Fig. S1A,B and S11, and Supplemental Table S1). Combination of alternating recognitions sites allows for higher degradation efficiency and reduces sequence repetitiveness. The two enzymes have recognition sites of lengths 27 and 18 bp, respectively, which even with some degeneracy allowed in the recognition site (Gimble and Wang 1996; Argast et al. 1998) makes their random occurrence in a transcriptome highly improbable. The two enzymes work at the same temperature and in the same buffer, so their digestion can be combined in a single step. A fraction of the synthesised carrier RNA is capped using Vaccinia Capping System (NEB) and mixed with uncapped carrier to achieve the desired capping percentage (Palazzo and Lee 2015).

**Figure 1.**
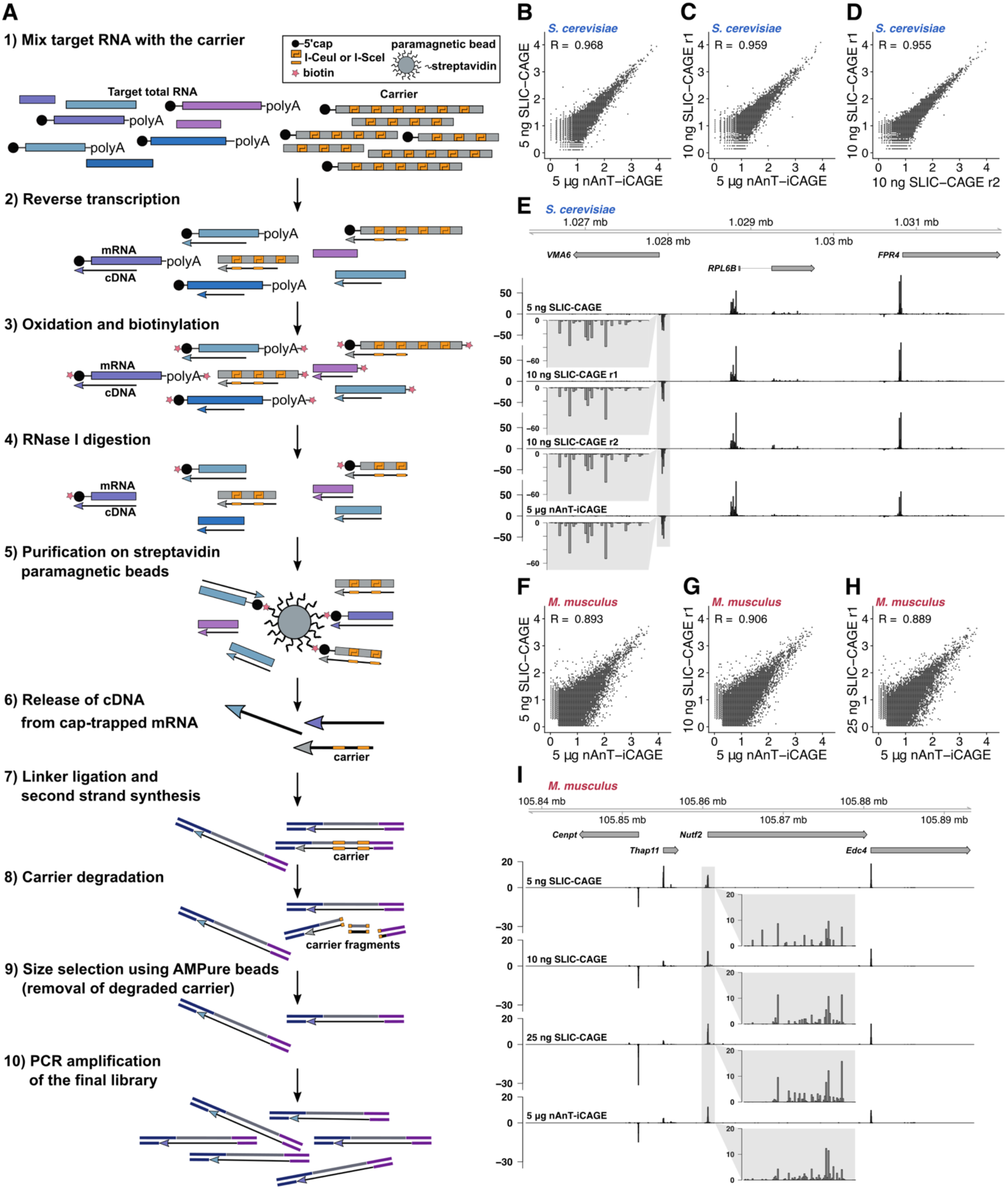
SLIC-CAGE development and assessment. (**A**) Schematics of the SLIC-CAGE approach. Target RNA of limited quantity is mixed with the carrier mix to get 5 μg of total RNA material. cDNA is synthesised through reverse transcription and cap is oxidized using sodium periodate. Oxidation allows attachment of biotin using biotin hydrazide. In addition to the cap structure, biotin gets attached to the mRNA’s 3’ end, as it is also oxidized using sodium periodate. To remove biotin from mRNA:cDNA hybrids with incompletely synthesized cDNA, and from mRNA’s 3’ ends, the samples are treated with RNase I. Complete cDNAs (cDNA that reached the 5’ end of mRNA), are selected by affinity purification on streptavidin magnetic beads (cap-trapping). cDNA is released from cap-trapped cDNA:mRNA hybrids and 5’-and 3’-linkers are ligated. The library molecules that originate from the carrier are degraded using I-Sce-I and I-Ceu-I homing endonucleases and the fragments removed using AMPure beads. The leftover library molecules are then PCR amplified to increase the amount of material for sequencing. (**B**-**C**) Pearson correlation at the CTSS level of nAnT-iCAGE and SLIC-CAGE libraries prepared from (**B**) 5 ng or (**C**) 10 ng of *S. cerevisiae* total RNA. (**D**) Pearson correlation at the CTSS level of SLIC-CAGE technical replicates prepared from 10 ng of *S. cerevisiae* total RNA. The axes in (B-D) show log10(TPM+1) values and the correlation was calculated on raw, non-log transformed data. (**E**) CTSS signal in example locus on chromosome 12 in SLIC-CAGE libraries prepared from 5 or 10 ng of *S. cerevisiae* total RNA, and in nAnT-iCAGE library prepared from standard 5 μg of total RNA. The inset grey boxes show a magnification of a tag cluster. (**F-H**) Pearson correlation at the CTSS level of nAnT-iCAGE and SLIC-CAGE libraries prepared from (**F**) 5 ng, (**G**) 10 ng or (**H**) 25 ng of *M. musculus* total RNA. The axes in (F-H) show log10(TPM+1) values and the correlation was calculated on raw non-log transformed data. (**I**) CTSS signal in example locus on chromosome 8 in SLIC-CAGE libraries prepared from 5, 10 or 25 ng of *M. musculus* total RNA, and in the reference nAnT-iCAGE library prepared from standard 5 μg of total RNA. The inset grey boxes show a magnification of a tag cluster and one bar represents a single CTSSs.

The percentage of capped RNAs in the carrier and its size distribution were optimised by performing the entire SLIC-CAGE protocol, starting by adding the synthetic carrier to the low-input sample to achieve a total of 5 μg of RNA material. To assess its performance, we compared its output with the nAnT-iCAGE library derived from 5 μg of total cellular RNA. We use nAnT-iCAGE as a reference as it is currently considered the most unbiased protocol for promoterome mapping (Murata et al. 2014), and because TSS identification by cap-trapper based technology has been experimentally validated (Carninci et al. 2006). To identify the optimal ratio of capped and uncapped carrier, as well as the length of the carrier RNAs, we tested the following carrier mixes: 1) carriers with lengths distributed between 0.3-1 kb *versus* homogenous 1 kb length carriers and 2) a mixture of capped and uncapped *versus* only capped carrier. We performed the SLIC-CAGE protocol outlined in Figure 1A, starting with 100 ng of total RNA isolated from *S. cerevisiae* supplemented with the various carrier mixes up to total 5 μg of RNA. We then compared the output with the nAnT-iCAGE library generated using 5 μg of total RNA (Supplemental Fig. S1C-M, Supplemental Tables S4 and S5, see Methods for more details). Removal of the carrier was performed by two rounds of degradation using homing endonucleases (I-SceI and I-CeuI, Supplemental Fig. S11) with a purification and a PCR amplification step between the rounds (see Methods for details of the SLIC-CAGE protocol). The presence of the carrier significantly improved the correlation of individual CAGE-supported TSSs (CTSSs, see Methods) between SLIC-CAGE and the reference nAnT-iCAGE library (Supplemental Fig. S1C-H). This effect was not observed when either only the capped carrier or no carrier was used (Supplemental Fig. S1C,G,H). The highest correlation and reproducibility was achieved by a carrier mix composed of 10% capped and 90% uncapped molecules of 0.3-1 kb length (Supplemental Fig. S1E,F, mix 2, Supplemental Tables S4 and S5). This mix was designed to closely mimic the composition of cellular total RNA (see Methods for more details). Other diagnostic criteria shown in Supplemental Fig. S1J-M confirm it as the optimal carrier choice.

### SLIC-CAGE allows genome-wide TSS identification from nanogram-scale samples

We set out to identify the lowest amount of total RNA that can be used to produce high quality CAGE libraries. To that end, we performed a SLIC-CAGE titration test with 1-100 ng of total *S. cerevisiae* RNA and compared this with nAnT-iCAGE library derived from 5 μg of total RNA. The high correlation of individual CTSSs between SLIC-CAGE and the reference nAnT-iCAGE library (Fig. 1B and C, Supplemental Fig. S2A) shows that genuine CTSSs are identified. Moreover, SLIC-CAGE libraries show high reproducibility (Fig. 1D). Figure 1E shows an example locus in the genome browser, demonstrating the high similarity of SLIC-CAGE and nAnT-iCAGE CTSS profiles in all high-quality datasets (*i.e*. datasets with high complexity, see below).

To confirm the general applicability of the SLIC-CAGE protocol, we performed a similar titration test using total RNA isolated from E14 mouse embryonic stem cells. The results obtained following sequencing of the libraries generated using 5, 10 or 25 ng of total RNA were highly correlated (Pearson correlation 0.9) with the reference nAnT-iCAGE derived library. The correlation did not improve further with increasing total starting RNA (Fig.1F-H, Supplemental Fig. S2C), again verifying the SLIC-CAGE protocol for nanogram-scale samples. The genome browser view (Fig. 1I) confirms the similarity of profiles on the individual CTSS level, although with minor differences in the library prepared from 5 ng of *M. musculus* total RNA due to lower complexity, as discussed in detail in the next section.

Analysis of library mapping efficiency demonstrated that selective degradation of the carrier is highly efficient. When only 1 ng of total RNA is used with a 5000-fold more carrier (5 μg), 25% of the sequenced reads are uniquely mapped to the target organism, while the rest corresponds to the leftover carrier (27%), short amplified linkers or multimappers, commonly discarded from TSS analyses (Supplemental Tables S10 and S11). This amount of leftover carrier is minor and does not significantly compromise sequencing depth (10% or less when 10 ng of total RNA are used). We expect that with additional rounds of degradation and purification, the leftover carrier could be further reduced, although with a risk of sample loss, and we found it unnecessary.

### Complexity and resolution of SLIC-CAGE libraries

Next, we wanted to explore library complexity and any potential inherent CTSS detection biases. As discussed above, both SLIC-CAGE and nAnT-iCAGE libraries are highly correlated at individual CTSSs, we thus set off in the next steps to examine the spatial clustering of these CTSSs and its features.

CTSSs in close vicinity reflect functionally equivalent transcripts and are generally clustered together and analysed as a single transcriptional unit termed a tag cluster (Haberle et al. 2015). The CTSS with the highest TPM value within a tag cluster is referred to as the *dominant* CTSS (see Methods). Specificity in capturing genuine TSSs can be assessed by examining the fraction of tag clusters that overlap with expected promoter regions. We identified a high percentage of SLIC-CAGE tag clusters that map to known promoter regions in both *S. cerevisiae* and *M. musculus* libraries irrespective of the total starting RNA, thus indicating the high specificity of these libraries (approximately 80%, at the same level as the reference nAnT-iCAGE protocol, Fig. 2A,E and Supplemental Fig. S2D,F).

**Figure 2.**
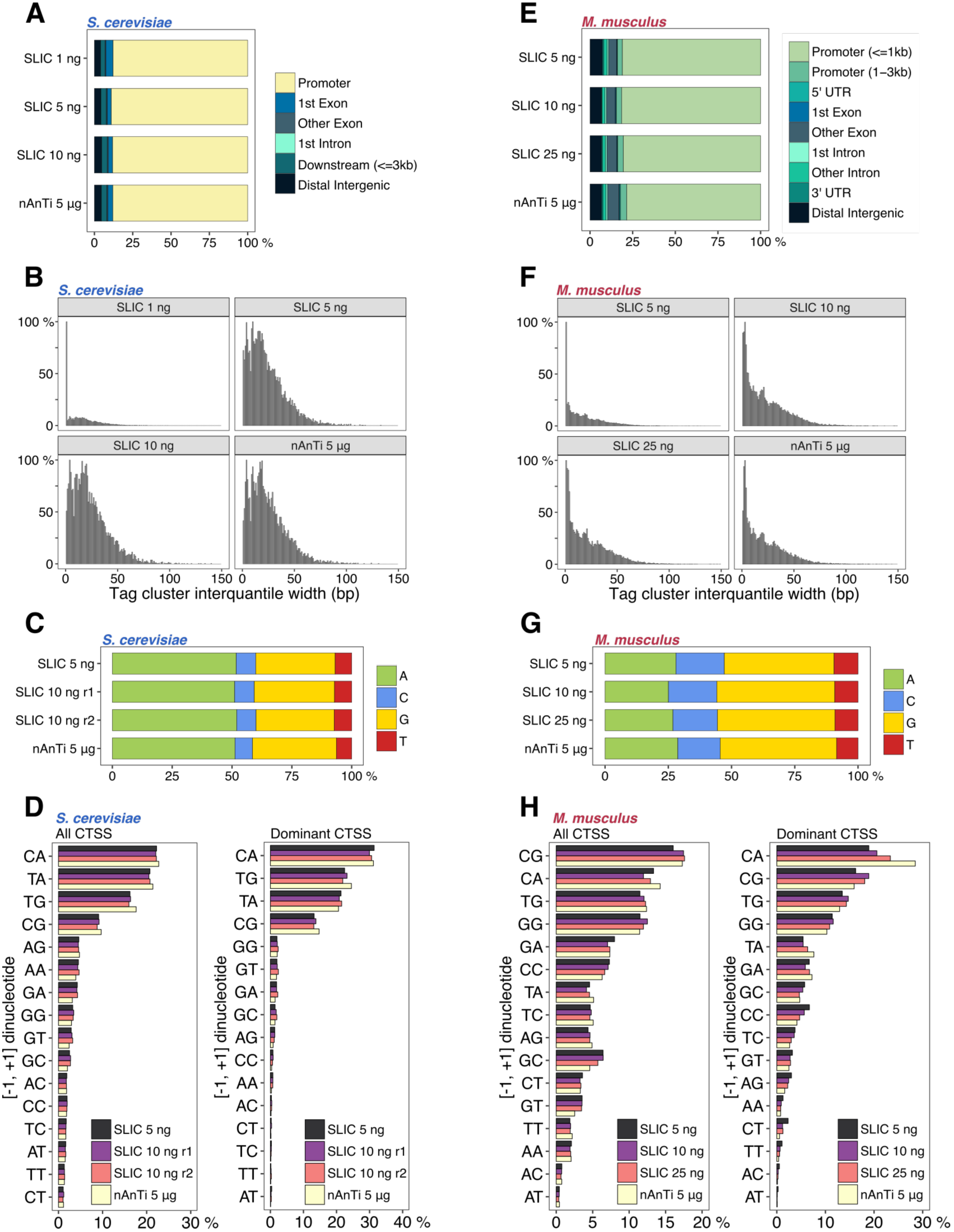
Identifying the lower limits of SLIC-CAGE libraries. **A**) Genomic locations of tag clusters identified in SLIC-CAGE libraries prepared from 1, 5 or 10 ng of *S. cerevisiae* total RNA *versus* the reference nAnT-iCAGE library. (**B**) Distribution of tag cluster interquantile widths in SLIC-CAGE libraries prepared from 1, 5 or 10 ng of *S. cerevisiae* total RNA and in the nAnT-iCAGE library. (**C**) Nucleotide composition of all CTSSs identified in SLIC-CAGE libraries prepared from 5 or 10 ng of *S. cerevisiae* total RNA and in the reference nAnT-iCAGE library. (**D**) Dinucleotide composition of all CTSSs (left panel) or dominant CTSSs (right panel) identified in SLIC-CAGE libraries prepared from 5 or 10 ng of *S. cerevisiae* total RNA and in the nAnT-iCAGE library. Both panels are ordered from the most to the least used dinucleotide in nAnT-iCAGE. (**E**) Genomic locations of tag clusters in SLIC-CAGE libraries prepared from 5, 10 or 25 ng of *M. musculus* total RNA and in the nAnT-iCAGE library. (**F**) Distribution of tag cluster interquantile widths in SLIC-CAGE libraries prepared from 5, 10 or 25 ng of *M. musculus* total RNA and the nAnT-iCAGE library. (**G**) Nucleotide composition of all CTSSs identified in SLIC-CAGE libraries prepared from 5, 10 or 25 ng of *M. musculus* total RNA or identified in the nAnT-iCAGE library. (**H**) Dinucleotide composition of all CTSSs (left panel) or dominant CTSSs (right panel) identified in SLIC-CAGE libraries prepared from 5, 10 or 25 ng of *M. musculus* total RNA or identified in the reference nAnT-iCAGE library. Both panels are ordered from the most to the least used dinucleotide in the reference nAnT-iCAGE.

In addition to determining the number of unique detected CTSSs and tag clusters, and their overlap with the reference library (Supplemental Table S12), complexity of CAGE-derived libraries can be assessed by comparing tag cluster widths. To robustly identify tag cluster widths, the interquantile widths (IQ-width) were calculated that span 10^th^ and the 90^th^ percentile (q0.1-q0.9) of the total tag cluster signal to exclude effects of extreme outlier CTSSs. The distribution of tag cluster IQ-widths serves as a good visual indicator of library complexity, as in low-complexity libraries, incomplete CTSS detection will lead to artificially sharp tag clusters. This low-complexity effect can be simulated by randomly subsampling a high-complexity nAnT-iCAGE library (see Supplemental Figure S3). IQ-width distribution of *S. cerevisiae* SLIC-CAGE tag clusters reveals that complexity of the reference nAnT-iCAGE library is recapitulated using as little as 5 ng of total RNA (Fig. 2B, Supplemental Fig. S4A). This result is substantiated with the number of unique CTSSs which corresponds to the number identified with nAnT-iCAGE (around 70% overlap between 5 ng SLIC-CAGE and nAnTi-iCAGE, and 90% overlap in tag cluster identification). Low-complexity with artificially sharper tag clusters is seen only with 1-2 ng of total RNA input (Fig. 2B, Supplemental Fig. S4A and Supplemental Table S12). A highly similar result is observed with *M. musculus* SLIC-CAGE libraries, although lower complexity is notable at 5 ng of total RNA (Fig. 2F and Supplemental Fig. S4B). This is in agreement with the lower number of unique CTSSs identified in 5 ng *M. musculus* SLIC-CAGE library compared to nAnT-iCAGE (Supplemental Table S12). We expect that an increase in sequencing depth would ultimately recapitulate the complexity of the reference dataset as higher coverage in *S. cerevisiae* facilitates higher complexity libraries with lower starting amount (5 ng).

To assess sensitivity and precision of SLIC-CAGE, we used standard RNA-seq receiver operating characteristic (ROC) curves with true CTSSs and tag clusters defined by the nAnT-iCAGE library, and show similar ratios of true and false positives when identifying CTSSs or tag clusters in high complexity libraries, regardless of total RNA input amount (Supplemental Figure S5 A and C).

We also assessed SLIC-CAGE derived CTSS features from *S. cerevisiae* and *M. musculus* and compared them with features extracted using the nAnT-iCAGE library as reference. First, nucleotide composition of all SLIC-CAGE-identified CTSSs reveals highly similar results to nAnT-iCAGE independent of the total input RNA (Fig. 2C,G and Supplemental Fig. S2G,I). Furthermore, the composition of [-1,+1] dinucleotide initiators (where the +1 nucleotide represents the identified CTSS) also showed a highly similar pattern to the reference nAnT-iCAGE dataset (Fig. 2D,H left panel, Supplemental Fig. S2J,L). SLIC-CAGE libraries identify CA, TA, TG and CG as the most preferred initiators, similar to preferred mammalian initiator sequences (Carninci et al. 2006).

Focusing only on the initiation patterns ([-1, +1] dinucleotide) of the dominant TSS (CTSSs with the highest TPM within each tag cluster) of each tag cluster facilitates estimating influence of PCR amplification on the distribution of tags within a tag cluster. Highly similar dinucleotide composition of dominant TSS initiators, independent of the amount of total RNA used, confirms that identification of the dominant TSSs is not obscured by PCR amplification (Fig. 2D,H right panel, Supplemental Fig. S2M,O). The identified preferred initiators are pyrimidine-purine dinucleotides CA, TG, TA (*S. cerevisiae*) or CA, CG, TG (*M. musculus*) in accordance with the Inr element (YR) (Burke and Kadonaga 1997; Haberle and Lenhard 2016). These results confirmed the utility of SLIC-CAGE in uncovering authentic transcription initiation patterns such as the well-established CA initiator.

Further, we analysed the distance between dominant TSSs identified in tag clusters common to each SLIC-CAGE and the reference nAnt-iCAGE sample and show: 1) the same dominant TSS is identified in 50-60% cases in high complexity libraries (>= 10 ng of total RNA); 2) 75-80% of identified dominant TSSs are within 10 bp distance of the nAnT-iCAGE-identified dominant TSS in high complexity libraries (Supplemental Figures S6 and S7, Supplemental Table 12). Some variability between the identified dominant TSSs is expected, even between technical nAnT-iCAGE replicates, especially in broad tag clusters with several TSSs of similar expression level (70% of dominant TSSs within 10 bp distance of nAnT-iCAGE technical replicates, Supplemental Table S13, Supplemental Figure S8)

As a final assessment of SLIC-CAGE performance, we analysed expression ratios per individual CTSS common to SLIC-CAGE and the reference nAnT-iCAGE (Supplemental Fig. S9 and S10 left panels) and present the ratios in a heatmap centred on the dominant CTSS identified by the reference nAnT-iCAGE library. This analysis can uncover any positional biases, if introduced by the SLIC-CAGE protocol. Patterns of signal in heatmaps (grouping upstream or downstream of the nAnTi-iCAGE-identified dominant CTSS) would signify positional bias and indicate non-random capturing of authentic TSSs. We also evaluated the positions and expression values of CTSSs identified in the nAnT-iCAGE but absent in SLIC-CAGE libraries (Supplemental Fig. S9 and S10, middle panels). We found there are no positional biases with regards to SLIC-CAGE-identified CTSSs and their expression values, independent of the total input RNA. As expected, a higher number of CTSSs identified in nAnT-iCAGE were absent from lower complexity *S. cerevisiae* SLIC-CAGE libraries derived from 1 and 2 ng total RNA (Supplemental Fig. S9A,B, middle panels). This was particularly evident in those CTSSs with expression values in the lower two quartiles (top two sections in each heatmap). Further, the CTSSs identified in both low-complexity SLIC-CAGE and nAnT-iCAGE exhibit higher TPM ratios, likely reflecting the effect of PCR amplification. On the other hand, we found that the SLIC-CAGE library derived from 5 ng of total RNA (Supplemental Fig. S9C) shows similar patterns as libraries derived from greater amounts of RNA (Supplemental Fig. S9D-H) or the library derived by PCR amplification of the nAnT-iCAGE library (Supplemental Fig. S9I).

Similar results were obtained when comparing *M. musculus* SLIC-CAGE libraries with their reference nAnT-iCAGE library (Supplemental Fig. S10) albeit with a twofold greater minimum starting RNA (10 ng) required for high-complexity libraries. Overall, these results show that SLIC-CAGE increases the sensitivity of the CAGE method 1000-fold over the current “gold standard” nAnT-iCAGE, without decrease in signal quality. This unparalleled sensitivity without loss of fidelity of start base capture positions SLIC-CAGE as a method of choice for unbiased identification of TSSs in low-input samples that were previously inaccessible to CAGE methodology.

### SLIC-CAGE generates superior quality libraries compared to existing low input methods

The current available method for low input samples, nanoCAGE, requires 50-500 ng of total cellular RNA and is very different from standard CAGE in its selection of capped RNAs (Plessy et al. 2010; Poulain et al. 2017). Whilst the gold standard verified CAGE protocols, *i.e.* nAnT-iCAGE relies on cap-trapper based selection of capped RNA, nanoCAGE uses the template switching property of the reverse transcriptase to selectively introduce a barcoded adapter only onto 5’ ends of capped RNA. The result are hybrid cDNA molecules with a specific nucleotide sequence added to the 5’ end of the capped RNA.

We first set off to compare nanoCAGE against nAnT-iCAGE. We carried out a nanoCAGE titration test using *S. cerevisiae* total RNA (5-500 ng) and compared the obtained libraries with the reference nAnT-iCAGE library. CTSSs identified in nanoCAGE libraries were poorly correlated (Pearson correlation 0.5-0.6) with the nAnT-iCAGE library, irrespective of the total RNA used (Fig. 3A-E and Supplemental Fig. S2B). Despite reduced similarity with nAnT-iCAGE, nanoCAGE libraries appeared reproducible (Fig. 3F). An example genome browser view also reveals significant differences in CTSS profiles between nanoCAGE and nAnT-iCAGE libraries (Fig. 3G). NanoCAGE systematically failed to capture all CTSSs identified with nAnT-iCAGE. In contrast, our SLIC-CAGE library derived from only 5 ng of total RNA accurately recapitulates the nAnT-iCAGE TSS profile shown in the same genomic region (Fig. 3G as Fig. 1E).

**Figure 3.**
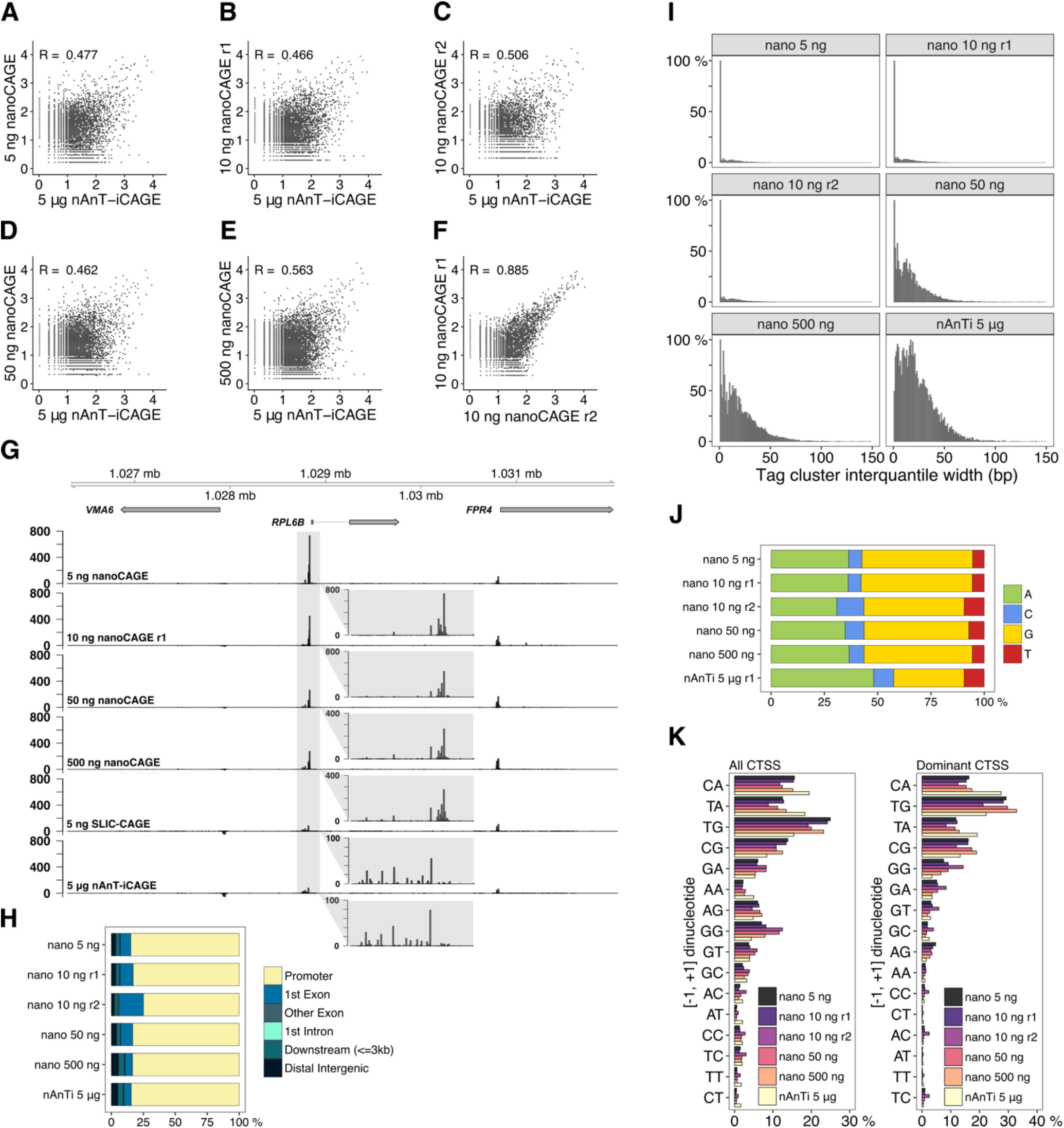
Comparison of nanoCAGE and the reference nAnT-iCAGE. (**A-E**) Pearson correlation of nAnT-iCAGE and nanoCAGE libraries prepared from (**A**) 5 ng, (**B, C**) 10 ng, (**D**) 50 ng or (**E**) 500 ng of *S. cerevisiae* total RNA. (**F**) Pearson correlation of nanoCAGE technical replicates prepared from 10 ng of *S. cerevisiae* total RNA. (**G**) CTSS signal in example locus on chromosome 12 in nanoCAGE libraries prepared from 5, 10, 50 or 500 ng, SLIC-CAGE library prepared from 5 ng, and the nAnT-iCAGE library prepared from 5 μg of *S. cerevisiae* total RNA (the same locus is shown in Fig. 1E). The inset grey boxes show a magnification of a tag cluster. Insets in nanoCAGE libraries have a different scale, as signal is skewed with PCR amplification. Different tag cluster is magnified compared to Fig. 1E, as nanoCAGE did not detect the upstream tag cluster on the minus strand (**H**) Genomic locations of tag clusters identified in nanoCAGE libraries prepared from 5-500 ng of *S. cerevisiae* total RNA and in the nAnT-iCAGE library (**I**) Distribution of tag cluster interquantile widths in nanoCAGE libraries prepared from 5-500 ng of *S. cerevisiae* total RNA *versus* the reference nAnT-iCAGE library. (**J**) Nucleotide composition of all CTSSs identified in nanoCAGE libraries prepared from 5-500 ng of *S. cerevisiae* total RNA or identified in the reference nAnT-iCAGE library. (**K**) Dinucleotide composition of all CTSSs (left panel) or dominant CTSSs (right panel) identified in nanoCAGE libraries prepared from 5-500 ng of *S. cerevisiae* total RNA or in the reference nAnT-iCAGE library. Both panels are ordered from the most to the least used dinucleotide in the reference nAnT-iCAGE.

Next, we investigated the tag clusters identified in each nanoCAGE library and showed that approximately 85% were indeed in expected promoter regions (Fig. 3H, and Supplemental Fig. S2E). The cluster overlap is highly similar to the reference nAnT-iCAGE library in all nanoCAGE libraries, independent of the amount of total RNA used. Therefore, nanoCAGE does not capture the full complexity of TSS usage but its specificity for promoter regions is not diminished.

To inspect the complexity of nanoCAGE libraries, we again compared tag cluster IQ-widths with the reference nAnT-iCAGE library (Fig. 3I and Supplemental Fig. S4C). An increase in the number of sharper tag clusters is observed at 1-50 ng of total input RNA. The IQ-width distributions show that nanoCAGE systematically produces lower-complexity libraries compared to nAnT-iCAGE and SLIC-CAGE. This result agrees well with the consistently lower number of unique CTSSs identified in nanoCAGE libraries and its low overlap with nAnT-iCAGE CTSSs (Supplemental Table S13).

To further assess performance of nanoCAGE against nAnT-iCAGE and SLIC-CAGE, we used standard RNA-seq receiver operating characteristic (ROC) curves with true CTSSs or tag clusters defined by the nAnT-iCAGE library (Supplemental Figure S5C). In comparison to SLIC-CAGE, nanoCAGE performs poorly - the ratio of true and false positives is low and highly dependent on total RNA input amount.

Nucleotide composition of nanoCAGE-identified robust CTSSs revealed a strong preference for G-containing CTSSs (Fig. 3J). This is specific to nanoCAGE libraries and independent of the starting total RNA amount. This observed G-preference is not an artefact caused by the extra C added complementary to the cap structure at the 5’end of cDNA during reverse transcription, as that is common to all CAGE protocols and corrected using the Bioconductor package CAGEr (Haberle et al. 2015). Lastly, to check if in nanoCAGE more than one G is added during reverse transcription, we counted the 5’end Gs flagged as a mismatch in the alignment and found that the amount of two consecutive mismatches was not significant (Supplemental Table S14).

We also analysed the distance between dominant TSSs identified in tag clusters common to each nanoCAGE and the reference nAnT-iCAGE sample and show that: 1) nanoCAGE captures only 30% of the dominant TSSs regardless of the RNA input quantity; 2) only about 60% of identified dominant TSSs are within 10 bp distance of the nAnT-iCAGE identified dominant TSS regardless of the RNA input (Supplemental Figure S8, Supplemental Table S13). Taken together, the results demonstrate that SLIC-CAGE strongly outperforms nanoCAGE in identification of true dominant TSSs.

The composition of [-1,+1] initiator dinucleotides revealed a severe depletion in identified CA and TA initiators, with the corresponding increase in G-containing initiators (TG and CG), in comparison with the reference nAnT-iCAGE dataset (Fig. 3K, left panel, Supplemental Fig. S2K). To assess the most robust CTSSs, we repeated the same analysis using only the dominant CTSSs in each tag cluster (Fig. 3K, right panel, Supplemental Fig. S2N) and the lack of CA and TA initiators was equally apparent. This property of nanoCAGE makes it unsuitable for the determination of dominant CTSSs and details of promoter architecture at base pair resolution.

To exclude the effects of CTSSs located in non-promoter regions and to assess if CTSS identification depends on expression levels, we divided tag clusters according to their genomic location or expression values (division into four expression quartiles per each library) and repeated the analysis (Supplemental Fig. S11 and S12). Since a similar pattern (depletion of CA and TA initiators) was observed irrespective of the genomic location or expression level, these results suggest that the nanoCAGE bias is caused by template switching of reverse transcriptase known to be sequence dependent and expected to preferentially capture capped RNA that starts with G (Zajac et al. 2013).

Finally, we analysed signal ratios of individual CTSSs identified in each nanoCAGE library and the reference nAnT-iCAGE (ratio of TPM values, Supplemental Fig. S13 left panels), and CTSSs not identified in nanoCAGE (Supplemental Fig S13 middle panels) similarly as described for SLIC-CAGE (see above). This analysis reveals that there are no position specific biases in nanoCAGE and that the biases are primarily caused by nucleotide composition of the capped RNA 5’ends. Further, it accentuates the inability of nanoCAGE to capture dominant CTSSs identified with the reference nAnT-iCAGE, even with higher amounts of starting material, compared to SLIC-CAGE (Supplemental Fig. S13F-H *vs* Supplemental Fig. S9A-H).

To ensure that the observed nanoCAGE biases are general and not specific to our library, we analysed and compared nAnT-iCAGE and nanoCAGE XL data recently produced on human K562 cell line (data from (Adiconis et al. 2018)). Our results were recapitulated with Adiconis et al datasets.

The nanoCAGE XL library exhibited low correlation with the nAnTi-iCAGE library at individual CTSSs and tag clusters expression level (Supplemental Figure S14A and B). Although distribution of interquantile widths suggests that libraries are not of low complexity (Supplemental Figure S14D), this may be a consequence of contamination with non-capped captured RNA, as nanoCAGE XL poorly captures promoter regions (Supplementary Figure S14E). In addition, only 7% of the dominant CTSSs identified in nanoCAGE XL libraries matched nAnT-iCAGE identified dominant CTSSs, while CTSS and initiator biases were even more prominent (Supplemental Figure S14C and G) than in our dataset.

### Using SLIC-CAGE to uncover promoter architecture

The dominant CTSS provides a structural reference point for the alignment of promoter sequences and thus facilitates the discovery of promoter-specific sequence features. High-quality data is necessary for the accurate identification of the dominant TSS within a tag cluster or promoter region. Sharp promoters, described by small IQ-widths, are typically defined by a fixed distance from a core promoter motif, such as a TATA-box or TATA-like element at -30 position (Ponjavic et al. 2006) upstream of the TSS, or by DPE motif at +28 to +32 (Kutach and Kadonaga 2000) in Drosophila. Broader vertebrate promoters, featuring multiple CTSS positions, are enriched for GC content and CpG island overlap and also exhibit precisely positioned +1 nucleosomes (Haberle and Lenhard 2016). Lower complexity libraries have an increased number of artificially sharp tag clusters (Fig. 2F) due to sparse CTSS identification. Although the identified CTSSs in lower-complexity SLIC-CAGE libraries are canonical, their association of sequence features may be obscured by artificially sharp tag clusters as they will group with true sharp clusters and dilute the signal derived from sequence features (see analysis with 5 ng sample below). To address this, we investigated the promoter architecture for known promoter features in E14 mouse embryonice stem cells (mESC) using SLIC-CAGE from 5 to 100 ng of total RNA.

We first assessed the presence of a TA dinucleotide around the -30 positions for all CTSSs identified by SLIC-CAGE for both 5 and 10 ng of input RNA. The dominant CTSSs were ordered by IQ-width of their corresponding tag cluster and extended to include 1 kb DNA sequence up-and downstream. The TA frequency is depicted in a heatmap in Fig. 4A for promoters ordered from sharp to broad for 10 ng of RNA and clearly recapitulates the patterns visible in the reference nAnT-iCAGE library (similarity of heatmaps is assessed by permutation testing, Supplemental Figure S15A). As expected, the sharpest tag clusters in libraries produced from 5 ng of total RNA have a weaker TA signal (Supplemental Fig. S16A, TA heatmap), as these are likely artificially sharp and not the canonical sharp promoters. A similar result is observed for enrichment of the canonical TATA-box element, the 10 ng library recapitulated the reference nAnT-iCAGE library whereas the 5 ng library shows a weaker enrichment (Fig. 4B and Supplemental Fig. S15B and S17C,D).

**Figure 4.**
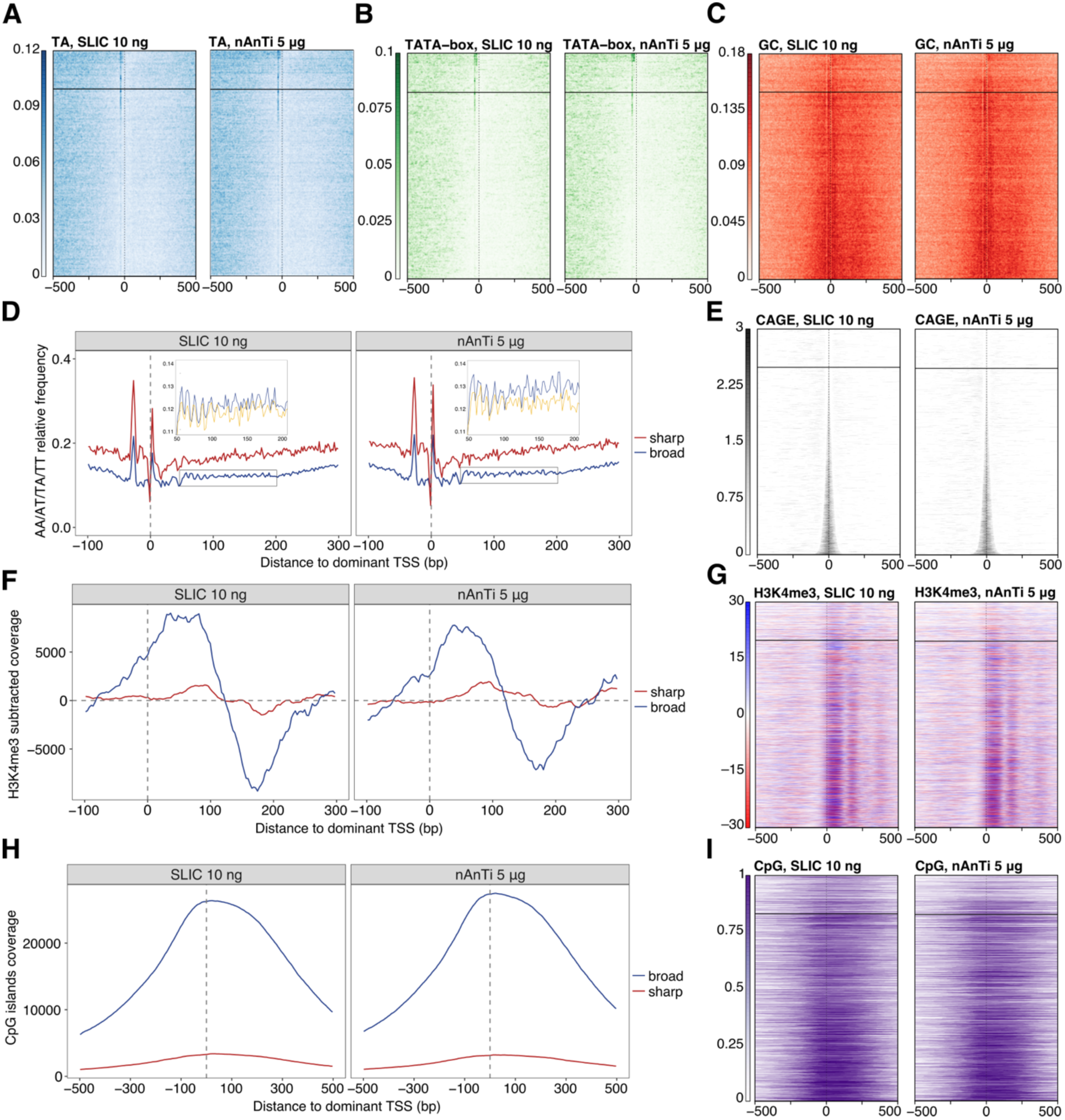
SLIC-CAGE is equivalent to nAnT-iCAGE for pattern discovery. Comparison of SLIC-CAGE derived from 10 ng and nAnT-iCAGE derived from 5 μg of *M. musculus* total RNA. In all heatmaps, promoters are centred at the dominant CTSS (dashed vertical line at 0) and ordered by tag cluster interquantile width with sharpest promoters on top and broadest on the bottom of each heatmap. The horizontal line separates sharp and broad promoters (empirical boundary for sharp promoters is set at interquantile width <= 3). (**A**) Comparison of TA dinucleotide density in the SLIC-CAGE (left) and nAnT-iCAGE library (right). As expected, sharp promoters have a strong TA enrichment, in line with the expected TATA-box in sharp promoters around the -30 position. (**B**) Comparison of TATA-box density in SLIC-CAGE (left, 23.6% of TCs have a TATA-box around -30 position) vs nAnT-iCAGE library (right, 24.1% of TCs have a TATA-box around -30 position). Promoter regions are scanned using a minimum of 80^th^ percentile match to the TATA-box position weight matrix (PWM). Sharp promoters exhibit a strong TATA-box signal as suggested in A). (**C**) Comparison of GC dinucleotide density in the SLIC-CAGE (left) and nAnT-iCAGE library (right). Broad promoters show a higher enrichment of GC dinucleotides across promoters, suggesting presence of CpG islands, as expected in broad promoters. (**D**) Average WW (AA/AT/TA/TT) dinucleotide frequency in sharp and broad promoters identified in SLIC-CAGE (left) or nAnT-iCAGE library (right). Inset shows a closer view on WW dinucleotide frequency (blue) overlain with the signal obtained when the sequences are aligned to a randomly chosen identified CTSS within broad promoters (yellow). 10 bp WW periodicity implies the presence of well-positioned +1 nucleosomes in broad promoters, in line with the current knowledge on broad promoters. (**E**) Tag cluster coverage heatmap of SLIC-CAGE (left) or nAnT-iCAGE library (right). (**F**) H3K4me3 relative coverage in sharp versus broad promoters identified in SLIC-CAGE (left) or nAnT-iCAGE (right). Signal enrichment in broad promoters indicates well positioned +1 nucleosomes, in line with the presence of WW periodicity in broad promoters. (**G**) H3K4me3 signal density across promoter regions centred on SLIC-CAGE or nAnT-iCAGE identified dominant CTSS. (**H**) Relative coverage of CpG islands across sharp and broad promoters, centred on dominant CTSS identified in SLIC-CAGE (left) or nAnT-iCAGE (right). These results are in agreement with GC-dinucleotide density signal, which is much stronger in broad promoters. (**I**) CpG islands coverage signal across promoter regions centred on dominant CTSS identified in SLIC-CAGE (left, 68.1% of TCs overlap with a CpG island) or nAnT-iCAGE (right, 64.4% of TCs overlap with a CpG island).

A GC-enrichment in the region between the dominant TSS and 250 bp downstream of it indicates positioning of the +1 nucleosomes and is expected to be highly localized in broad promoters. This feature is again recapitulated by the 10 ng RNA input library (Fig. 4C, Supplemental Fig. S15C and S16) (Consortium et al. 2014; Haberle et al. 2014; Haberle and Lenhard 2016). Furthermore, rotational positioning of the +1 nucleosomes are associated with WW periodicity (AA/AT/TA/TT dinucleotides) lined up with the dominant TSS. Next, we examined WW dinucleotide density separately for sharp and broad promoters identified by SLIC-CAGE and the reference nAnT-iCAGE library (Fig. 4D). A strong 10.5 bp periodicity of WW dinucleotides downstream of the dominant TSS was observed in SLIC-CAGE libraries derived from 10 ng of *M. musculus* total RNA and corresponded to the phasing observed with the reference nAnT-iCAGE library (Fig. 4D and Supplemental Fig. S17B). This can only be observed across promoters if the dominant TSS is accurately identified and therefore it reflects the quality of the libraries. To confirm that WW dinucleotide periodicity reflects +1 nucleosome positioning in broad promoters, we assessed H3K4me3 data downloaded from ENCODE (Fig. 4F and G, Supplemental Fig. S16, H3K4me3 heatmap, Supplemental Fig. S15D). H3K4me3 subtracted coverage reflects the well-positioned +1 nucleosome broad promoters (Fig. 4F,G) and localizes with WW periodicity specific for broad promoters (Fig. 4D). These results are in agreement with previously identified nucleosome positioning preferences (Segal et al. 2006).

As a final validation of SLIC-CAGE promoters, we assessed CpG island density separately in sharp and broad promoters (Fig. 4H and I, Supplemental Fig. S15E and S16). We observed a higher density of CpG islands in SLIC-CAGE broad promoters, which corresponds to nAnT-iCAGE broad promoters and agrees with the expected association of broad promoters and CpG islands (Carninci et al. 2006; Haberle and Lenhard 2016). These results demonstrate the utility of SLIC-CAGE libraries derived from nanogram-scale samples in promoter architecture discovery, alongside the gold standard nAnT-iCAGE libraries.

### Uncovering the TSS landscape of mouse primordial germ cells using SLIC-CAGE

Transcriptome, epigenome and methylome changes occurring during primordial germ cell development have been thoroughly studied (Hajkova 2011; Yamaguchi et al. 2013; Hill et al. 2018). However, the total number of cells and the total amount of RNA obtained per embryo are severely limited. As a result, the underlying regulatory changes at the level of promoter activity and TSS usage have not been addressed, primarily due to lack of adequate low-input methodology. We applied SLIC-CAGE to mouse primordial germ cells embryonic (E) day 11.5 using approximately 10 ng of total RNA obtained from 5000-6000 cells isolated per litter (7-8 embryos) and provide its first promoterome/TSS landscape.

To validate PGC E11.5 SLIC-CAGE library, we compared CAGE-derived gene expression levels to a published PGC E11.5 RNA-seq dataset (Yamaguchi et al. 2013) and found high correlation (Spearman correlation coefficient 0.81, Figure 5A upper-left panel). The correlation between E11.5 PGC SLIC-CAGE and E11.5 PGC RNA-seq expression is significantly and reproducibly higher than between E11.5 PGC SLIC-CAGE and E14 mESC RNA-seq expression, although mESC E14 and PGC E11.5 have similar TSS/promoter landscapes (Figure 5A, lower-left and lower-right panels). High similarities of interquantile width distributions (Figure 5B), genomic locations of identified tag clusters (Figure 5C), CTSS and dominant CTSS dinucleotide distributions (Figure 5D and E) between SLIC-CAGE PGC E11.5 and nAnTi-CAGE mESC E14 libraries validate the high quality of the PGC E11.5 SLIC-CAGE dataset. Furthermore, we observed canonical promoter types such as sharp and broad promoters used in the PGC E11.5 stage. We also detected classification associated sequence characteristics, as sharp tag clusters/promoters were associated with the presence of TATA-box (23.7% of TCs), while broad tag clusters/promoters overlapped with CpG islands (63% of TCs, Figure 5F).

**Figure 5.**
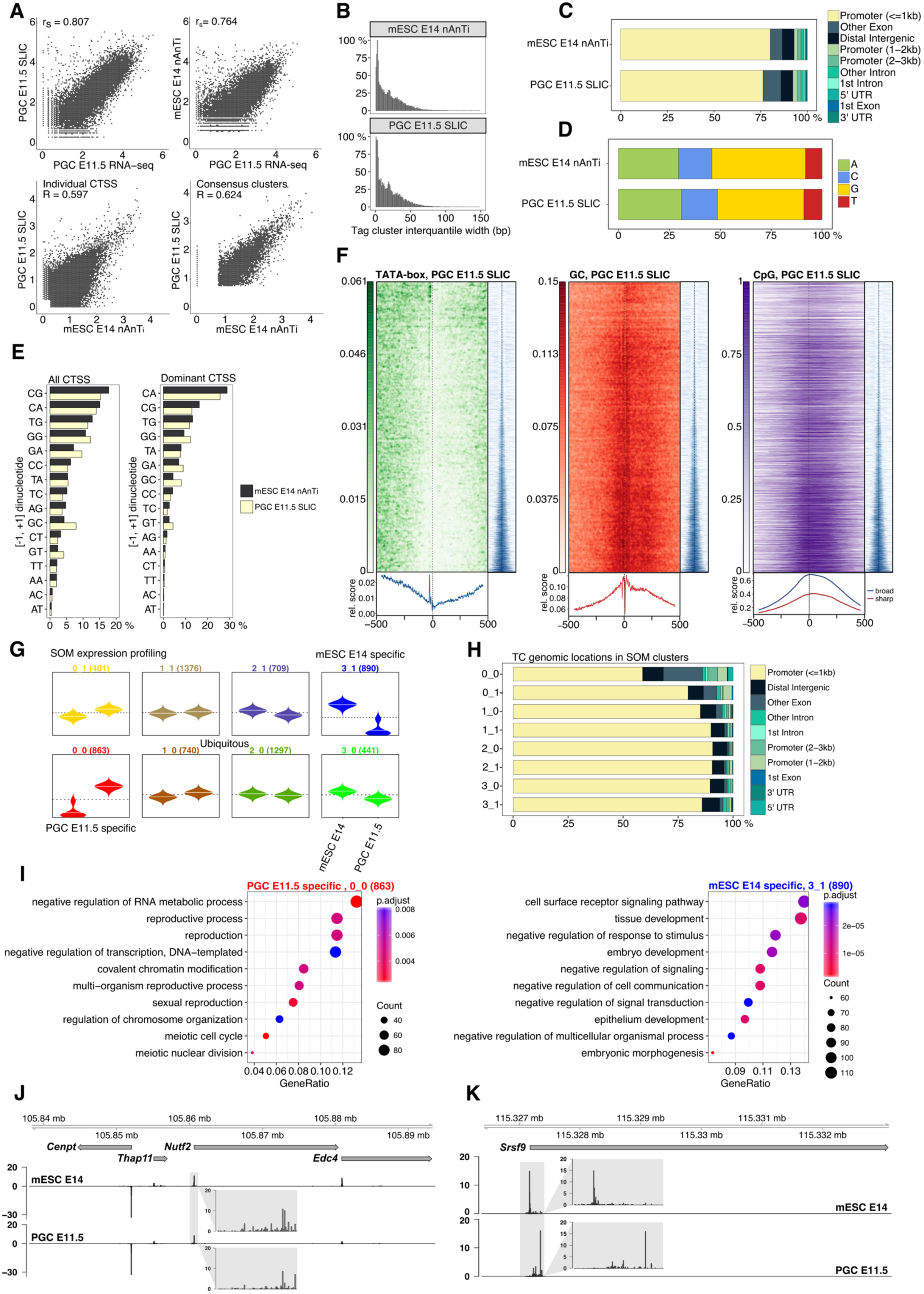
TSS landscape of primordial germ cell E11.5 stage. (**A**) Spearman correlation of SLIC-CAGE PGC E11.5 data with PGC E11.5 RNA-seq data (upper left) or nAnT-iCAGE mESC E14 data with PGC E11.5 RNA-seq data (upper right panel). Pearson correlation of SLIC-CAGE PGC E11.5 and nAnT-iCAGE mESC E14 datasets on individual CTSS level (lower left) or consensus tag cluster/promoter level (lower right panel). Comparison of nAnT-iCAGE mESC E14 and SLIC-CAGE PGC E11.5 libraries: (**B**) distribution of tag cluster interquantile widths; (**C**) genomic locations of tag clusters (**D**) nucleotide composition of all CTSSs; (**E**) dinucleotide composition of all CTSSs (left panel) or dominant CTSSs (right panel). Both panels are ordered from the most to the least used dinucleotide in the mESC E14 library. (**F**) TATA-box, GC dinucleotide and CpG island density in PGC E11.5 data. In all heatmaps, promoters are centred at the dominant CTSS (dashed vertical line at 0). Promoter regions are scanned using a minimum of 80^th^ percentile match to the TATA-box PWM. The signal metaplot is shown below each heatmap, and a tag cluster IQ-width coverage (in blue) shows ordering in the pattern heatmap from sharp to broad tag clusters/promoters (200 bp window centred on dominant TSS). (**G**) Expression profiles obtained by SOM clustering of tag-clusters/promoters. Each box represents one cluster, left beanplots represent mESC E14 and right beanplots represent PGC E11.5. The horizontal line denotes the mean expression level in each cluster. (**H)** Genomic locations of tag cluster in each SOM class (SOM-classes are shown on y-axis). (**I**) Biological process GO analysis in PGC E11.5 specific SOM class 0_0 (left) and mESC E14 specific class 3_1 (right). (**J**) CTSS signal in example locus on chromosome 8 (left panel, same as in Fig.1I) and (**K**) CTSS signal in *Srsf9* promoter region (chr5) exhibiting TSS switching in PGC E11.5 compared to mESC E14. The inset grey boxes show magnification of tag clusters.

Similar results were obtained with an independent biological replicate of PGC E11.5, for which we used approximately 10-15 ng of highly degraded total RNA (Supplemental Fig. S18A). Somewhat lower correlation of replicates and lower mapping to promoter regions is observed due to use of degraded RNA, however the data shows no bias in CTSS composition and allows capturing of canonical promoter features (Supplemental Fig. S18B-G). This proves that the SLIC-CAGE protocol can be used to generate unbiased TSS landscapes even from degraded samples, which is more often required when the samples are hard to obtain.

We also used the paired-end information on random reverse priming to collapse PCR duplicates in the replicate data – 47% of uniquely mapped reads are kept upon deduplication. We show that deduplicated data highly correlates with the data prior to deduplication on both CTSSs and tag cluster levels, causing no changes in interquantile-widths distribution (Supplemental Figure S19).

Using self-organising maps (SOM) expression profiling, we identified differentially and ubiquitously expressed tag clusters in PGC E11.5 stage vs mESC E14 cell line (Figure 5G, H). Biological process GO analysis of the PGC E11.5-specific SOM cluster 0_0 revealed enrichment of reproduction and meiosis-related terms (Figure 5I, left panel), while mESC E14 specific SOM cluster 3_1 showed tissue and embryo development terms (Figure 5I, right panel). In line with the recently discovered set of genes crucial for normal gametogenesis (45 germline reprogramming-responsive genes or GRRs) (Hill et al. 2018), we found nine GRR genes in the PGC E11.5 specific 0_0 SOM class (Rad51c, Dazl, Slc25a31, Hormad1, 1700018B24Rik, Fkbp6, Stk31, Asz1 and Taf9b), three GRR genes in the PGC E11.5 specific 0_1 SOM class (Mael, Sycp and Pnldc) and two in the ubiquitously expressed class 2_0 (D1Pas1 and Hsf2bp). Classes specific to mESC E14 cells or other ubiquitous classes did not contain any GRR genes.

Finally, we identified TSSs differentially used between mESC E14 cell line and PGC E11.5 stage. While the TSS landscapes are highly similar (Figure 5A, lower-left panel and Figure 5J left panel), we identified several genes with differential TSS usage within the same promoter region (Figure 5K, right panel). This may reflect differential preinitiation machinery, known to differ in gametogenesis (Goodrich and Tjian 2010), but also may cause alternative transcript/protein isoforms, change translational efficiency due to 5’UTR variation or alter differential transcript stability (Tamarkin-Ben-Harush et al. 2017; Leppek et al. 2018).

## DISCUSSION

We have developed SLIC-CAGE, an unbiased cap-trapper based CAGE protocol optimized for promoterome discovery from as little as 5-10 ng of isolated total RNA (approximately 10^3^ cells, RIN >= 7, as generally recommended for CAGE techniques (Takahashi et al. 2012)). SLIC-CAGE may also be used on low-quality RNA, however an increase in the amount of starting material may be required for high quality libraries. We show that SLIC-CAGE libraries are of equivalent quality and complexity as nAnT-iCAGE libraries derived from 500-1000-fold more material (5 μg of total RNA, approximately 10^6^ cells). SLIC-CAGE extends the nAnT-iCAGE protocol through addition of the degradable carrier to the target RNA material of limited availability. Since the best CAGE protocol is not amenable to downscaling, the idea behind the carrier is to increase the amount of material to permit highly specific cap-trapper based purification of target RNA polymerase II transcripts and to minimize material loss in many protocol steps.

We designed the carrier to have a similar size distribution and fraction of capped molecules as the total cellular RNA, to effectively saturate non-specific adsorption sites on all surfaces and matrices used throughout the protocol. In the final stage of SLIC-CAGE, the carrier molecules are selectively degraded using homing endonucleases, while the intact target library is amplified and sequenced. SLIC-CAGE, equally as nAnT-iCAGE (Murata et al. 2014), permits for paired-end sequencing and linking TSSs to transcript architecture. In addition, paired-end data contains information on random priming in reverse transcription, which may be used as a UMI to collapse identical read pairs as PCR duplicates.

We have shown that SLIC-CAGE is superior in sensitivity, resolution and absence of bias to the only other low-input CAGE technology, nanoCAGE, which relies on template switching during the cDNA synthesis (Plessy et al. 2010). Although the amount of starting material is significantly reduced, the lowest input limit for nanoCAGE is 50 ng of total RNA which may require up to 30 PCR cycles (Poulain et al. 2017). We directly compared performances of SLIC-CAGE and nanoCAGE in titration tests and demonstrated that: 1) higher complexity libraries are achieved with significantly lower input: SLIC-CAGE requires 5-10 ng, while nanoCAGE requires 50 ng of total RNA; 2) nanoCAGE does not recapitulate the complexity of the nAnT-iCAGE libraries even with the highest recommended amount of RNA (500 ng), in comparison SLIC-CAGE captures the full complexity when 5-10 ng are used; 3) nanoCAGE preferentially captures G-starting capped mRNAs, while SLIC-CAGE does not have 5’mRNA nucleotide dependent biases; 4) Biases in nanoCAGE libraries are independent of the total RNA amount used, and inherent to the template switching step. Our results are in agreement with a recent study that demonstrated low performance of nanoCAGE and other template-switching technologies in capturing true transcription initiation events, compared to cap-trapper based CAGE technologies (Adiconis et al. 2018). However, the effects of template-switching bias are presumably reduced when used for RNA-seq purposes and gene expression profiling. Overall, cap-trapper based CAGE methods outperform RAMPAGE, STRT, NanoCAGE-XL and Oligo-capping (Adiconis et al. 2018).

Importantly, with the carrier approach to minimize the target sample loss, SLIC-CAGE protocol requires less PCR amplification cycles – 15-18 cycles for 10-1 ng of total RNA as input. This is advantageous as smaller number of PCR cycles avoids amplification biases and the fraction of observed duplicate reads. Although, nanoCAGE takes advantage of unique molecular identifiers to remove PCR duplicates, in our experience, synthesis of truly random UMI is problematic and subject to variability, thereby obscuring its use.

A different carrier-based approach has recently been applied to down-scale chromatin-precipitation based methods – favoured amplification recovery via protection ChIP-seq (FARP-ChIP-seq) (Zheng et al. 2015). FARP-ChIP-seq relies on a designed biotinylated synthetic DNA carrier, mixed with chromatin of interest prior to ChIP-seq library preparation. Amplification of the synthetic DNA carrier is prevented using specific blocker oligonucleotides. These blocker oligonucleotides are: 1) complementary to the biotin-DNA carrier; 2) carry phosphorothioate modification of the first three nucleotides at the 5’end for resistance to exonuclease activity of the polymerase; 3) carry a 3’end 3-carbon spacer to inhibit extension by PCR. The blocker strategy can achieve a 99% reduction in amplification of the biotin-DNA, which if applied instead of our degradable carrier would leave much more carrier to sequence (starting SLIC-CAGE with 1 ng of total RNA and 5 μg of the carrier, 27% of the carrier is left in the final library, which is more than a 10000-fold reduction, Supplemental Table S11). While also being more costly and time consuming, this approach could be combined with selective degradation of the SLIC-CAGE carrier when near-complete removal of carrier is required thereby increasing sequencing depth to allow higher complexity libraries from even lower input amounts.

Finally, we show that SLIC-CAGE is applicable to low cell number samples by obtaining the TSS landscape of mouse PGC E11.5 stage. In comparison with mESC E14, we show that PGC E11.5 have highly similar features and canonical promoter signatures. We also identify genes specific to PGC E11.5 stage, further validating the high quality of the PGC E11.5 TSS atlas. Although the TSS landscape in shared promoter regions is similar between mESC E14 and PGC E11.5 stage, we uncover TSS switching events. Identification of TSS switching events and biological follow-up studies will lead to a higher understanding of its functional consequences.

We anticipate that SLIC-CAGE will prove to be invaluable for in-depth and high-resolution promoter analysis of rare cell types, including early embryonic developmental stages or embryonic tissue from a wide range of model organisms, which has so far been inaccessible to the method. With its low material requirement (5-10 ng of total RNA), SLIC-CAGE can also be applied on isolated nascent RNA, to provide an unbiased promoterome with high positional and temporal resolution. Lastly, as bidirectional capped RNA is a signature feature of active enhancers (Andersson et al. 2014), deeply sequenced SLIC-CAGE libraries can be used to identify active enhancers in rare cell types. The principle of the degradable carrier can also be easily extended to other protocols where the required amount of RNA or DNA is limiting.

## METHODS

### Preparation of the carrier RNA molecules

DNA template (1kb length) for preparation of the carrier by *in vitro* transcription was synthesized and cloned into pJ241 plasmid (service by DNA 2.0, Supplemental Fig. S1, Supplemental Table S1) to produce the carrier plasmid. The template encompasses the gene that serves as the carrier, embedded with restriction sites for I-SceI and I-CeuI to allow degradation in the final steps of the library preparation. The templates for *in vitro* transcription were prepared by PCR amplification using the unique forward primer (PCR_GN5_f1, Supplemental Table S2) which introduces the T7 promoter followed by five random nucleotides, and the reverse primer which determines the total length of the carrier template and introduces six random nucleotides at the 3’end (Supplemental Table S2, PCR_N6_r1-r10).

The PCR reaction to produce the carrier templates was composed of 0.2 ng μl^-1^ carrier plasmid, 1 μM primers (each), 0.02 U μl^-1^ Phusion High-Fidelity DNA Polymerase (Thermo Fisher Scientific) and 0.2 mM dNTP in 1 x Phusion HF Buffer (final concentrations). The cycling conditions are presented in Supplemental Table S3. Produced carrier templates (lengths 1034-386 bp) were gel-purified to remove non-specific products.

Carrier RNA was *in vitro* transcribed using HiScribe™ T7 High Yield RNA Synthesis Kit (NEB) according to manufacturer’s instructions and purified using RNeasy Mini kit (Qiagen). A portion of carrier RNAs was capped using Vaccinia Capping System and purified using RNeasy Mini kit (Qiagen). The capping efficiency was estimated using RNA 5′Polyphosphatase and Terminator ™5′-Phosphate-Dependent Exonuclease, as only uncapped RNAs are dephosphorylated and degraded, while capped RNA’s are protected.

Several carrier combinations were tested in SLIC-CAGE (Supplemental Table S4 and S5) and the final carrier used in SLIC-CAGE was comprised of 90% uncapped carrier and 10% capped carrier, both of varying length (Supplemental Table S5).

We note that the carrier RNA mix can be prepared within two days in large quantities for multiple library preparations and frozen at -80°C until use.

### Sample collection and nucleic acid extraction

*S. cerevisiae* BY4741 strain was grown in YPD media, and when the cells reached the exponential phase, collection was done by centrifugation. The cell material was stored at -80 degrees C prior to RNA isolation.

*S. cerevisiae* total RNA was extracted from using the standard hot-phenol procedure. The extracted RNA was additionally purified using the Qiagen RNeasy kit (clean-up protocol, according to manufacturer’s instructions). Isolated RNA was quantified using NanoDrop1000 and the quality of RNA assessed on the bioanalyzer (Agilent). The RNA samples were of high quality (RIN > 9).

Mouse E14 cells were grown in in N2B27 (recipe below) supplemented with the inhibitors LIF (Millipore, ESG1107), CHIR99021 (Cambridge Bioscience, SM13-10) and PD-0325901 (Caltag-Medsystems Limited, SYN-1059). Cells were detached by trypsinization, collected by spinning down and frozen at -80 degrees C prior to RNA isolation.

Total RNA from mouse embryonic stem cells (E14 cell line) was extracted using Qiagen RNeasy kit (according to manufacturer’s instructions). Isolated RNA was quantified using NanoDrop1000 and the quality of RNA assessed on the bioanalyzer (Agilent). The RNA samples were of high quality (RIN > 9).

### SLIC-CAGE library preparation

For the standard cap analysis of gene expression, the latest nAnT-iCAGE protocol was followed (Murata et al. 2014). In the SLIC-CAGE variant, the carrier was mixed with the RNA of interest, to the total amount of 5 μg, *e.g.* 10 ng of RNA of interest were mixed with 4990 ng of carrier mix and subjected to reverse transcription as in the nAnT-iCAGE protocol (Murata et al. 2014). Further library preparation steps were followed as described in Murata *et al* 2014(Murata et al. 2014) with several exceptions: 1) samples were pooled only prior to sequencing, to allow individual quality control steps; 2) samples were never completely dried using the centrifugal concentrator and then redissolved as in nAnT-iCAGE, instead the leftover volume was monitored to avoid complete drying and adjusted with water to achieve the required volume; 3) After the final AMPure purification in the nAnT-iCAGE protocol, each sample was concentrated using the centrifugal concentrator, and its volume adjusted to 15 μl, out of which 1 μl was used for quality control on the Agilent Bioanalyzer HS DNA chip.

Steps regarding degradation of the carrier in SLIC-CAGE libraries are schematically presented in Supplemental Fig. S20.

To degrade the carrier, 14 μl of sample was mixed with I-SceI (5 U) and I-CeuI (5 U) in 1 x CutSmart buffer (NEB) and incubated at 37 °C for 3 h. The enzymes were heat inactivated at 65°C for 20 min, and the samples purified using AMPure XP beads (1.8 x AMPure XP volume per reaction volume, as described in Murata *et al* (Murata et al. 2014)). The libraries were eluted in 42 μl of water and concentrated to 20 μl using the centrifugal concentrator.

A qPCR control was then performed to determine the suitable number of PCR cycles for library amplification and assess the amount of the leftover carrier. The primers designed to amplify the whole library are complementary to 5’ and 3’ linker regions, while the primers used to selectively amplify just the carrier are complementary to the 5’end of the carrier (common to all carrier molecules) and the 3’linker (common to all molecules in the library, see Supplemental Table S6 for primer sequences). qPCR reactions were performed using KAPA SYBR FAST qPCR kit using 1 μl of the sample and 0.1 μM primers (final concentration), in 10 μl total volume using PCR cycle conditions presented in the Supplemental Table S7.

The number of cycles for PCR amplification of the library corresponded to the C_t_ value obtained with the primers that amplify the whole library (adapter_f1 and adapter_r1, Supplemental Table S6). PCR amplification of the library was then performed using KAPA HiFi HS ReadyMix, with 0.1 μM primers (adapter_f1 and adapter_r1, Supplemental Table S6) and 18 μl of sample in a total volume of 100 μl. The cycling programme is presented in the Supplemental Table S8 and the final number of cycles used to amplify the libraries in Supplemental Table S9. Amplified samples were purified using AMPure XP beads (1.8 x volume ratio of the beads to the sample), eluted with 42 μl of water and concentrated using centrifugal concentrator to 14 μl.

A second round of carrier degradation was then performed as described for the first round. The samples were purified using AMPure XP beads (stringent 1:1 AMPure XP to sample volume ratio to exclude primer dimers and short fragments), eluted with 42 μl of water and concentrated to 12 μl using centrifugal concentrator. The combination of 1^st^ round of carrier degradation followed by PCR amplification, AMPure XP purification and the 2^nd^ round of carrier degradation is necessary to avoid substantial sample loss that leads to low-complexity libraries.

Each sample was then individually assessed for fragment size distribution using an HS DNA chip (Bioanalyzer, Agilent). If short fragments were present in the library (< 300 bp, see Supplemental Fig. S21), another round of size selection was performed using a stringed volume ratio of AMPure XP beads to the sample – 0.8 x (volume of each sample was prior to purification adjusted with water to 30 μl). The samples were eluted in 42 μl of water and concentrated to 12 μl using centrifugal concentrator. Fragment size distribution was again checked using an HS DNA chip (Bioanalyzer, Agilent), to ensure removal of the short fragments.

Finally, the amount of leftover carrier was estimated using qPCR as described above after the 1^st^ round of carrier degradation. The expected C_t_ in qPCR using adapter_f1 and adapter_r1 is 12-13 or 23-30 using carrier_f1 and adapter_r1 primer pairs (Supplemental Table S6) when the starting total RNA amount is 100-1 ng.

The libraries were sequenced on MiSeq (*S. cerevisiae*) or HiSeq2500 (*M. musculus*) Illumina platforms in single-end 50 base-pair mode (Genomics Facility, MRC, LMS),

### NanoCAGE library preparation

*S. cerevisiae* nanoCAGE libraries were prepared as described in the latest protocol version by Poulain *et al* 2017(Poulain et al. 2017). Briefly, 5, 10, 25, 50 or 500 ng of *S. cerevisiae* total RNA was reversely transcribed in the presence of corresponding template switching oligonucleotides (Supplemental Table S15) followed by AMPure purification. One 500 ng replicate was pre-treated with exonuclease to test if rRNA removal has any effects on the quality of the final library.

The number of PCR-cycles for semi-suppressive PCR was determined by qPCR as described in Poulain *et al* 2017(Supplemental Table S9). Samples were AMPure purified after amplification, and the concentration of each sample determined using Picogreen.

2 ng of each sample were pooled prior tagmentation and 0.5 ng of the pool was used in tagmentation. The sample was AMPure purified and quantified using Picogreen prior to MiSeq sequencing in single-end 50 base-pair mode (Genomics Facility, MRC, LMS).

### PGC E11.5 isolation and SLIC-CAGE library preparation

E11.5 PGCs were isolated from embryos obtained from a 129Sv female and GOF18ΔPE-EGFP (Yoshimizu et al. 1999) male cross. Briefly, genital ridges from one litter (7-8 embryos) were dissected out and digested at 37°C for 3 min using TrypLE Express (Thermo Fisher Scientific). Enzymatic digestion was neutralized with DMEM/F-12 (Gibco) supplemented with 15% fetal bovine serum (Gibco) followed by manual dissociation by pipetting. The cells were spun down by centrifugation and resuspended in 0.1% BSA PBS. GFP-positive cells were isolated using an Aria Fusion (BD Bioscience) flow cytometer and sorted into ice-cold PBS. Total RNA was isolated from approximately 5000-6000 E11.5 PGCs per litter using DNA/RNA Duet Kit miniprep kit (Zymo Research, USA). The SLIC-CAGE library was then prepared by mixing the obtained ~ 10 ng of PGC E11.5 total RNA (measured by Agilent 2100 Bioanalyzer) with 5 μg of the carrier mix and processed as described in SLIC-CAGE library preparation section.

### Processing of CAGE tags: nAnT-iCAGE, SLIC-CAGE or nanoCAGE

Sequenced CAGE tags (50 bp) were mapped to a reference *S. cerevisiae* genome (sacCer3 assembly) or *M. musculus* genome (mm10 assembly) using Bowtie2 (Langmead and Salzberg 2012) with default parameters that allow zero mismatches per seed sequence (default 22 nucleotides). Sequenced nanoCAGE libraries were trimmed prior to mapping to remove the linker and UMI region (15 bp from the 5’end were trimmed). FASTQ files from Adiconis et al (Adiconis et al. 2018) K562 nanoCAGE XL and nAnT-iCAGE libraries (replicate 1) were obtained from the SRA database (SRR6006247 and SRR6006235). As all nAnT-iCAGE libraries produced in that study were highly correlated, we chose to use only one replicate to match nanoCAGE XL (only one replicate was produced for nanoCAGE XL). Only read1 was used from both libraries and mapped using Bowtie2 (as described above) to hg19.

Only uniquely mapped reads were used in downstream analysis within R graphical and statistical computing environment (http://www.R-project.org/) using Bioconductor packages (http://www.bioconductor.org/) and custom scripts. The mapped reads were sorted and imported into R as bam files using CAGEr (Haberle et al. 2015). The additional G nucleotide at the 5’end of the reads, if added through template free activity of the reverse transcriptase, was resolved within CAGEr’s standard workflow designed to remove G’s that do not map to the genome: 1) if the first nucleotide is G and a mismatch, *i.e.* it does not map to the genome, it is removed from the read; 2) if the first nucleotide is G and it matches, it is retained or removed according to the percentage of mismatched G.

Sample replicates and reads from different lanes were merged prior to the final analysis as presented in Supplemental Material (Supplemental Tables S16-18). All unique 5’ends represent CAGE tag-supported TSSs (CTSSs), and the number of tags within each CTSS represents expression levels. Raw tag counts were normalized using a referent power-law distribution to a total of 10^6^ tags, resulting in normalized tags per million (TPMs) (Balwierz et al. 2009).

Deduplication of mouse PGC E11.5 replicate 2 was done using Clumpify from BBMap suite. Information from paired-end reads was utilised to collapse PCR duplicates. Since random TCT-N_6_ primer is used in reverse transcription, identical read pairs are expected to originate from PCR duplicates.

### Clustering of CTSSs into tag clusters and identification of dominant CTSS

CTSSs that pass the threshold of 1 TPM in at least one of the samples were clustered using a distance-based method implemented in the CAGEr package with a maximum allowed distance of 20 bp between the neighbouring CTSS.

For each tag cluster, a cumulative distribution of signal was calculated, and the boundaries of the tag cluster calculated using the 10^th^ and 90^th^ percentile of its signal. The distance between these boundaries represents the interquantile width of a tag cluster. The CTSS with the highest TPM value within a tag cluster is identified as the dominant CTSS (as implemented within CAGEr).

### Genomic locations of tag clusters

Tag clusters were annotated with their corresponding genomic locations using the ChIPseeker package (Yu et al. 2015). In *S. cerevisiae* libraries, promoters were defined as 1 kb windows centred on Ensembl (Aken et al. 2016) annotated transcriptions start sites (annotations imported from SGD) and in *M. musculus* libraries, promoters were defined as <= 1 kb or 1-3 kb from the UCSC annotated transcription start site.

### Nucleotide and dinucleotide composition of CTSSs

CTSSs from each library were filtered prior to analysis to include only CTSS with at least 1 TPM. In each library the number of A, C, G or T-containing CTSS was counted, divided by the total number of filtered CTSSs and converted to a percentage. The same analysis was performed using only dominant TSS (identified using the CAGEr package as a CTSS with highest expression within a tag cluster).

For dinucleotide analysis, identified filtered CTSSs were extended to include one upstream nucleotide ([-1, +1] dinucleotides where +1 represents the identified CTSS) and the same analysis as described above repeated for 16 possible dinucleotides.

### ROC curves

To assess accuracy of TSS identification for SLIC-CAGE and nanoCAGE libraries, we used nAnT-iCAGE libraries to define the set of true CTSSs and tag clusters. A true positive CTSS or a tag cluster corresponds to the CTSS or tag cluster in the nAnT-iCAGE library, while a false positive CTSS or a tag cluster exists only in the nanoCAGE or SLIC-CAGE library. ROC curves were generated in dependence of the CTSS or tag cluster TPM threshold in nanoCAGE or SLIC-CAGE libraries.

### Dinucleotide pattern analysis in *M. musculus* libraries

Heatmaps Bioconductor package (Perry M (18). heatmaps: Flexible Heatmaps for Functional Genomics and Sequence Features. R package version 1.2.0) was used to visualize dinucleotide patterns (TA and GC) across sequences centred on the dominant TSS. Sequences were ordered by interquantile width of the containing tag cluster, with the sharpest on top and broadest tag cluster on the bottom of the heatmap. Raw data with the exact matching for TA or GC was smoothed prior to plotting using kernel smoothing within the heatmaps package. Each heatmap was divided into two sections based on tag cluster’s IQ-widths. Empirical boundary (Supplemental Fig. S17A) was set to separate sharp (IQ-width <= 3 bp) and broad (IQ-width > 3) tag clusters identified in *M. musculus* libraries. The horizontal line/boundary was implemented using heatmaps options to partition heatmaps/rows of an image. Similarity of patterns between libraries was assessed by calculating the Jaccard distance between vectorized image matrices of smoothed heatmaps. Background similarity was assessed through calculation of Jaccard distance between vectorized image matrices of column-randomized, smoothed heatmaps. Column randomization was performed 10000 times, and the distribution of Jaccard distances calculated per each permutation was plotted and compared to the true Jaccard distance.

### TATA-box motif analysis in *M. musculus* libraries

SeqPattern package was used to scan the sequences for the occurrence of the TATA-box motif using a threshold of 80^th^ percentile match to the TATA-box PWM (imported from the seqPattern package). We further smoothed the obtained results using the kernel smoothing (heatmaps package) and plotted the results with sequences ordered by interquantile width of the containing tag cluster (sharpest on top and broadest on bottom of the tag cluster) and centred on the dominant TSS. The horizontal line in each heatmap represents the empirical boundary that separates sharp (IQ-width <= 3) and broad tag clusters (IQ-width > 3). Similarity between heatmaps was assessed as described above.

TATA-box metaplots (average signal/profile) were produced separately for sharp and broad tag clusters (see definition above). SeqPattern was used for scanning sequences using TATA-box PWM to identify 80% matches. The results were converted to the average signal using the heatmaps package with a 2 bp bin size. The final data was plotted using the ggplot2 package (Wickham 2009).

### Nucleosome positioning signal in in *M. musculus* libraries – WW periodicity

WW dinucleotide (AA/AT/TA/TT) occurrence (average relative signal) was obtained using the heatmaps package separately for sharp and broad tag clusters (see definition above). A 2 bp bin size was used and the sequences were centred on the dominant TSS. As a control for the importance of centring the sequences on the dominant TSS, WW dinucleotide (AA/AT/TA/TT) occurrence was obtained as an average relative signal from sequences where each sequence is centred on a randomly chosen CTSS within a tag cluster. The final data was plotted using the ggplot2 package (Wickham 2009).

### H3K4me3 signal around *M. musculus* tag clusters

H3K4me3 data for E14 cell line, mapped to mm10 was downloaded from ENCODE experiment ENCSR000CGO. Bam files for two replicates (accession numbers ENCFF997CAQ and ENCFF425ZMWO) were merged using samtools (Li et al. 2009) and the merged bam file was imported to R environment using the rtracklayer package (Lawrence et al. 2009)

H3K4me3 coverage was calculated separately for reads mapping to minus or plus strand and minus strand reads subtracted from plus strand reads to get the subtracted H3K4me3 coverage.

Subtracted H3K4me3 coverage was visualized using heatmaps package centred on the dominant TSSs with sequences ordered by IQ-width of the containing tag clusters (sharpest on top, and broadest at the bottom of the heatmap). Each heatmap was divided into two sections based on tag cluster’s IQ-widths as described above. Similarity between heatmaps was assessed as described above.

H3K4me3 coverage metaplots were produced separately for sharp and broad tag clusters (see definition above, only strongly supported dominant CTSSs with at least 5 TPM were used) using heatmaps package with a 3 bp bin size The final data was plotted using the ggplot2 package (Wickham 2009).

### *M. musculus* tag cluster overlap with CpG islands

The CpG island track for mm10 was downloaded from the UCSC Genome Browser. Overlap with *M. musculus* tag clusters was visualized as a coverage heatmap using heatmaps package, centred on the dominant TSS with sequences ordered by IQ-width of the containing tag clusters (sharpest on top, and broadest at the bottom of the heatmap). Each heatmap was divided into two sections based on tag cluster’s IQ-widths as described above.

CpG coverage metaplots were produced separately for sharp and broad tag clusters (see definition above) using heatmaps package with a 3 bp bin size. The final data was plotted using the ggplot2 package (Wickham 2009).

## DATA ACCESS

All data generated in this study – nAnT-iCAGE, SLIC-CAGE and nanoCAGE libraries have been submitted to ArrayExpress (https://www.ebi.ac.uk/arrayexpress/) under accession numbers E-MTAB-6519 and E-MTAB-7056 (mouse PGC E11.5 data).

## ACKNOWLEDGMENTS

This work was supported by The Wellcome Trust grant (106954) awarded to B.L. and F.M., MRC Core Funding (MC-A652-5QA10) and BBSRC Responsive Mode Grant (BB/R002703/1) awarded to H.G.L. N.C. was supported by an EMBO Long-Term Fellowship (EMBO ALTF 1279-2016), B.L. is supported by the Medical Research Council UK (MC UP 1102/1). H.G.L acknowledges support from the National Institute for Health Research (NIHR) Imperial Biomedical Research Centre (BRC). We thank Vedran Franke, Leonie Roos, Elena Pahita, Alexander Nash and Dunja Vucenovic for critical reading of the manuscript. We also thank Laurence Game and the MRC LMS sequencing facility for support.

## AUTHOR CONTRIBUTIONS

N.C. and B.L. conceived the study. P.H, F.M and P.C. advised development of the method. N.C. performed all experiments and computational analyses. H.G.L., M.B. and P.H. provided mouse RNA. N.C. and B.L. wrote the manuscript with input from all authors.

